# The H3.3 chaperone Hira complex orchestrates oocyte developmental competence

**DOI:** 10.1101/2020.05.25.114124

**Authors:** Rowena Smith, Zongliang Jiang, Andrej Susor, Hao Ming, Janet Tait, Marco Conti, Chih-Jen Lin

**Author notes:** equal contribution (Rowena Smith, Zongliang Jiang).

## Abstract

Reproductive success relies on a healthy oocyte competent for fertilisation and capable of sustaining early embryo development. By the end of oogenesis, the oocyte is characterised by a transcriptionally silenced state, but the significance of this state and how it is achieved remains poorly understood. Histone H3.3, one of the H3 variants, has unique functions in chromatin structure and gene expression that are cell cycle-independent. We report here a comprehensive characterisation of the roles of the subunits of the Hira complex (i.e. Hira, Cabin1 and Ubn1), which is primarily responsible for H3.3 deposition during mouse oocyte development. Loss-of-function of any component of the Hira complex led to early embryogenesis failure. Transcriptome and nascent RNA analyses revealed that mutant oocytes fail to silence global transcription. Hira complex mutants are unable to establish the H3K4me3 and H3K9me3 repressive marks, resulting in aberrant chromatin accessibility. Among the misregulated genes in mutant oocytes is Zscan4, a 2-cell specific gene that is involved in zygote genome activation. Overexpression of Zscan4 recapitulates the phenotypes of Hira mutants, illustrating that temporal and spatial expression of Zscan4 is fine-tuned at the oocyte-to-embryo transition. Thus, the H3.3 chaperone Hira complex has a maternal effect function in oocyte developmental competence and early embryogenesis by modulating chromatin condensation and transcriptional quiescence.

## Introduction

Oocytes are key orchestrators of fertilisation, initiation of zygotic genome activation and pre-implantation development ^1^. This potential is defined as developmental competence. Oocytes defective in developmental competence are a major cause of infertility, a medical condition which affects one in six couples. One hallmark feature of a competent oocyte is chromatin condensation. During the final stage of oocyte growth, oocytes with less condensed chromatin, termed non-surrounded nucleolus (NSN) oocytes, gradually transition to more condensed, surrounded nucleolus (SN) oocytes ^2^. Transcriptional silencing occurs around this time ^3^. However, the molecular components involved have not been defined and the primary underlying mechanism remains elusive ^4^.

Oocytes enter a protracted meiotic arrest in prophase I and maintain it for a long time, for years in humans, remarkably retaining their competence and genomic integrity. It would then be logical to hypothesise that replication-independent mechanisms of chromatin regulation may be indispensable to maintain the quality of oocytes throughout this prolonged meiotic arrest.

Histones are the fundamental blocks of chromatin that regulate both chromatin compactness and gene transcriptional processes. In addition to canonical histones, histone variants provide additional layers of regulatory mechanisms in a replication-independent manner. H3.3 is the primary replication-independent histone variant and has been shown to play prominent roles during critical stages of reproduction, including fertilisation ^5, 6^ and early embryonic development ^5, 7^. However, potential roles of H3.3 in oocyte transcriptional quiescence remain unexplored.

The key chaperone molecules required for H3.3 incorporation into chromatin form the Hira complex, which comprises Hira, Ubn1, and Cabin1 subunits ^8, 9^. In this study, we comprehensively dissect the roles of the Hira complex (Hira, Cabin1, and Ubn1) in establishment and maintainance of developmental competence in mouse oocytes. We demonstrate that the Hira complex is critical for oocyte developmental competence by stabilising a repressive epigenomic status that maintains transcriptional quiescence in preparation for embryogenesis.

## Results

### The maternal Hira Complex is essential for oocyte developmental competence

To investigate the maternal role of Hira complex during the oocyte-to-embryo transition, we knocked down Ubn1 in oocytes using an antisense morpholino approach and created new conditional knockout mouse lines targeting Hira and Cabin1, respectively (Fig. 1A). Firstly, we adopted the maternal knockdown approach we developed ^10^ to inhibit Ubn1 translation in fully-grown, germinal vesicle (GV) oocytes using antisense morpholino microinjection (Fig. S1A). We then rederived oocyte specific knockout mouse lines for Hira and Cabin1. We achieved this by crossing a transgenic Zp3-Cre mouse line with a conditional allele of Hira to deplete exon 6-7 (hereafter referred to as ZH in this study; the previously published Hira knockout was for exon 4; Fig. S1B) and similarly the Cabin1 mutant line was created by depletion of exon 6 (hereafter referred to as CabZ in this study; Fig. S1C).

**Figure 1.**
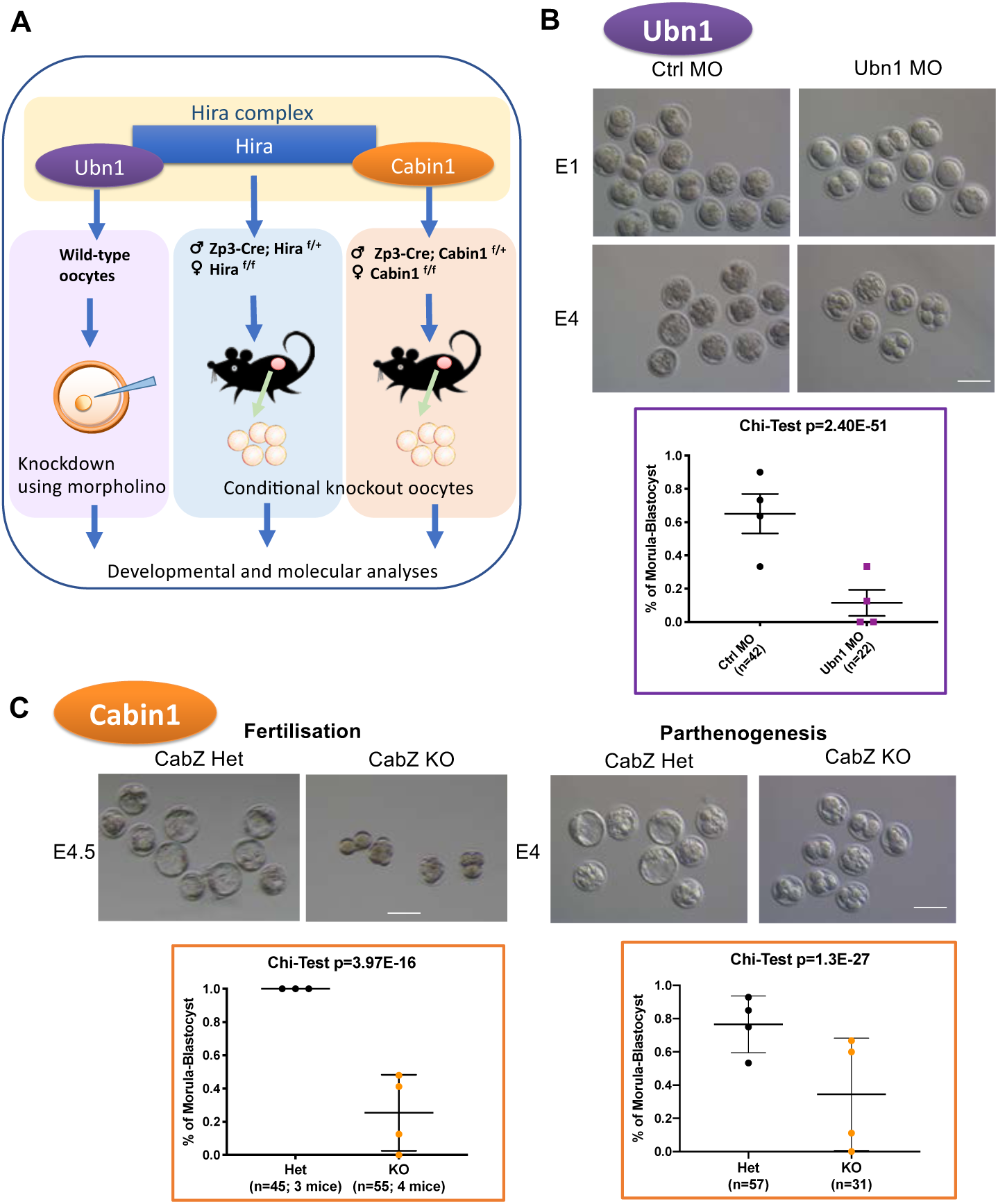
The maternal Hira complex is essential for pre-implantation development. (A) Oocyte specific loss-of-function approaches for each subunit of Hira complex. Ubn1 was knocked down by microinjection of morpholino antisense oligonucleotides. Conditional Hira and Cabin1 knockout mice were generated by crossing a Zp3-Cre line with Hira^flox/flox^ and Cabin1^flox/flox^ lines, respectively. (B) Ubn1 is required for pre-implantation development of parthenogenetic embryos. Developmental progression of control morpholino injected (Ctrl MO) and Ubn1 morpholino injected (Ubn1 MO) embryos at embryonic day1 (E1) and embryonic day 4 (E4). Left panel: Representative brightfield images; right panel: quantification. Scale bar: 100μm. (C) Both fertilised and parthenogenetically activated Cabin1 mutant oocytes arrested during preimplantation development. Left panel: Morula-to-blastocyst developmental potential of fertilised mutant Cabin1 embryos is impaired. Right panel: Morula-to-blastocyst developmental potential of parthenogenetically activated mutant Cabin1 embryos is impaired. Scale bar: 80μm.

To investigate whether maternal Ubn1 is involved in oocyte developmental competence, we compared the developmental potential of oocytes injected with control antisense morpholino (Ctrl MO) to those injected with Ubn1 antisense morpholino (Ubn1 MO). After microinjection into GV oocytes respectively, we performed *in vitro* maturation. Oocytes that matured to the MII stage were then used for parthenogenesis and pre-implantation development to the morula-to-blastocyst stage (Fig. S1A). Oocyte depletion of Ubn1 impaired maturation (55%, compared to Ctrl MO group 65%; p=1.7E-04; Fig. S1D). Importantly, it induced a significant decrease in cleavage to 2-cell (47%, compared to Ctrl MO group 78%; p=3.4E-06; Fig. S1E) as well as development to morula-to-blastocyst stage (11%, compared to Ctrl MO group 65%; p=2.4E-51; Fig. 1B).

Our new ZH mutant mice, with the exon 6-7 depletion, display the same phenotypes as those previously reported ^10^. Mice are infertile and form abnormal zygotes. CabZ mutant mice also generate abnormal zygotes (the post-fertilisation roles of Cabin1 and Ubn1 will be reported in a separate study, manuscript in preparation). To examine the developmental potential of the oocytes depleted of Cabin1 of the Hira complex, we flushed fertilised zygotes from CabZ heterozygous controls (CabZ Het) and mutants (CabZ KO), monitored their developmental outcomes, and compared the rate of development to morula-to-blastocyst stage embryos after *in vitro* culture. CabZ KO embryos had significantly reduced ability to form morula-to-blastocyst stage embryos (25%, compared to CabZ Het group of 100%; p=4E-16; Fig. 1C), the majority arrested at the 2-4 cell stages (Fig. 1C), which suggests a defect in zygotic genome activation ^11, 12^.

To further examine the maternal Cabin1 contribution to development, we performed parthenogenetic activation on *in vitro* matured MII oocytes from CabZ Het and CabZ KO mice. Parthenogenetic CabZ KO embryos failed to develop to morula-to-blastocyst stage and arrested at the 2-4 cell stage (34%, compared to CabZ Het group of 77%; p=1.3E-27; Fig.1C). These results firmly support that the developmental competence of mutant Cabin1 oocytes is compromised as assessed both with *in vivo* fertilisation or with parthenogenetic activation and therefore without contribution from the paternal genome.

Next, we asked whether the depletion of H3.3 in GV wild type oocytes would cause developmental arrest. Using the same methodology as Ubn1 knockdown, we knocked-down H3.3 in wild type oocytes. In agreement with the loss-of-function of H3.3 chaperone Hira complex (Hira, Cabin1 and Ubn1), knocking down H3.3 in oocytes resulted in the developmental arrest of parthenogenetic embryos by significantly impairing the formation of both 2-cell (74%, compared to Ctrl MO group of 91%; p=5.7E-5; Fig. S1F) and later morula-to-blastocyst stage embryos (6%, compared to Ctrl MO group of 87%; p=1.5E-124; Fig. S1F).

Taken together, these results demonstrate that loss of function of any one of the subunits of H3.3-Hira complex in oocytes induces failure of developmental progression through pre-implantation. Thus, the Hira complex is responsible for the acquisition of developmental competence in mouse oocytes, the corresponding genes functioning as maternal-effect genes. To confirm the timing and efficiency of loss-of-function is comparable in the same genetic models, we focused on characterising the phenotypes of maternal ZH or CabZ mutant oocytes (see below).

### Hira and Cabin1 pause expression of a cohort of embryogenetic genes in the oocyte

To gain insight into the cause of the embryonic arrest, we first investigated the transcriptional programmes in the oocytes with a disrupted Hira complex. The Zp3-Cre conditional approach achieves depletion of genes of interest until the formation of fully-grown oocytes at the end of oogenesis. To identify the genes and pathways affected by absence of Hira or Cabin1 in oocytes and precisely detect any earlier signs of aberrant transcriptomes, rather than collecting later stages oocytes (e.g. MII), we performed RNA sequencing (RNA-seq) in GV stage oocytes derived from both ZH and CabZ lines. A total of 1,462 and 207 genes were significantly differentially expressed (FDR adjusted p-value < 0.05; fold change > 2) in Hira and Cabin1 mutants compared to WT, respectively (Fig. 2A). Hira appeared to have a more profound impact on the transcriptome than Cabin1 (888 of upregulated and 574 downregulated transcripts in the ZH mutants; 154 upregulated and 53 downregulated transcripts in the CabZ mutants; Fig. 2A). Nevertheless, we observed that there is a noteworthy similarity in affected genes between ZH and CabZ mutants compared to WT oocytes. A greater percentage of the 57.5% of upregulated genes in CabZ mutants overlap with upregulated genes in ZH mutants; also 26.4% of CabZ differentially expressed (DE) downregulated genes overlapped with ZH downregulated genes (Fig. 2B). This finding of co-regulating genes by Hira and Cabin1 is consistent with observations in HeLa cells where separate knockdown of HIRA and CABIN1 ^9^ produced similar effects. These data also suggest that both subunits of Hira complex are equally important for the transcription outputs relevant for oocyte quality.

**Figure 2.**
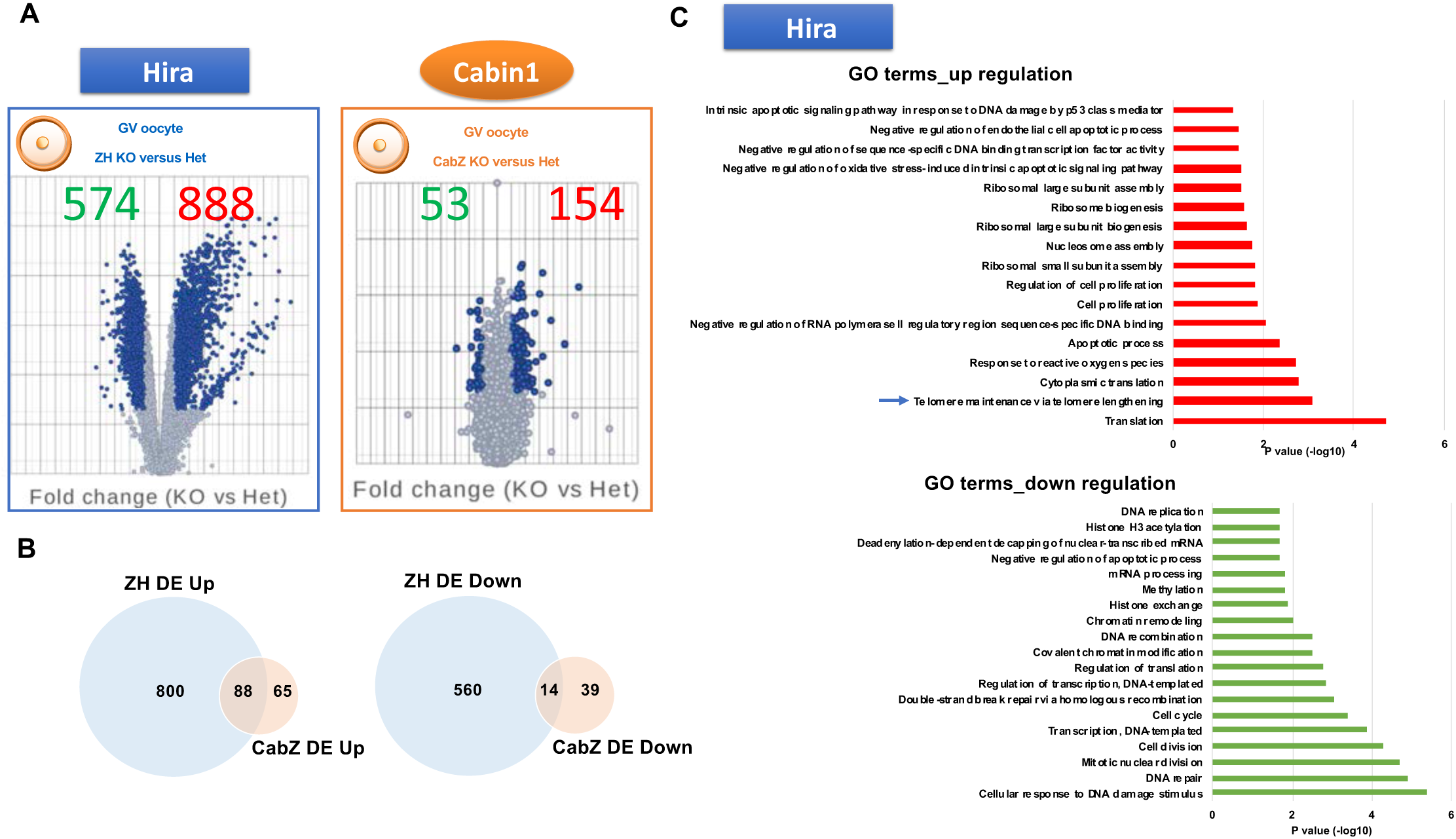
Loss-of-Hira complex impairs global transcription in GV oocytes. (A) Volcano plots showed differentially expressed genes of Hira and Cabin1 mutant GV oocytes respectively. Left panel: Differentially expressed genes of Hira mutant (ZH KO) compared to Hira heterozygous control (ZH Het) GV oocytes; right panel: Differentially expressed genes of Cabin1 mutant (CabZ KO) compared to Cabin1 heterozygous control (CabZ Het) GV oocytes. (B) Venn diagrams revealed the subsets of co-regulated genes dysregulated in Hira and Cabin1 mutant oocytes. 57.5% of Cabin1 mutant (CabZ KO) upregulated genes overlapped with Hira mutants (ZH KO). 26.4% of Cabin1 mutant (CabZ KO) downregulated genes overlapped with Hira mutants (ZH KO). (C) Top GO terms from up- and down-regulated genes of Hira mutant GV oocytes.

We noted that in general both ZH (888 versus 574) and CabZ (154 versus 53) inactivation causes more of an increase than a decrease in transcriptions in GV oocytes (Fig. 2A). This finding prompted us test whether there is a deficit of silencing global transcription in oocytes depleted of Hira complex. To exclude the possibility that the dysregulated genes were inherited from growing oocytes, we compared our datasets to a dataset generated by Ma et al ^13^, which extensively compared the transcriptomes of NSN (developmentally incompetent) to SN (developmentally competent) oocytes. Notably, both ZH and CabZ regulated genes revealed minimal overlapping with NSN oocytes (Fig. S2A), documenting that the overall dysregulation of the genes of ZH and CabZ mutants is not due to a developmental arrest at an immature oocyte stage.

We focused on identifying a cohort of genes that are regulated by Hira complex and that may be directly associated with oocyte developmental competence. Gene Ontology (GO) analyses showed that the up-regulated genes following Hira deletion are involved in translation, apoptotic process, and ribosomal biogenesis (Fig. 2C). Conversely, the down-regulated genes are involved in DNA damage and repair, transcription, and chromatin remodelling (Fig. 2C). Notably, among top-ranked GO terms from up-regulated genes was an enrichment of a set of telomere related genes (Fig. 2C). We further compared our data with 2-cell specific gene datasets generated by Wu et al ^14^ (additional analyses by Percharde et al ^15^, Fig. S2B), and found an overlap with 2-cell specific genes with both ZH and CabZ regulated genes (e.g. *Ddit3* and *Gm5039* and qRT-PCR validation in Fig. S2B). This camparison indicates that the depletion of Hira complex in oocytes resulted in the “derepression” of embryonic associated genes at the improper stages of development. The results suggest that the Hira complex may exert a repressive role to prevent precocious embryonic gene expression.

### Loss-of-Hira complex caused derepression of Zscan4 which correlated to the failure of cessation of transcription

We chose to focus on *Zscan4* as a candidate gene since it is involved in telomere maintenance. Also, it is one of the 2-cell specific genes, critical for regulating pluripotency/totipotency ^16, 17^. Zscan4 transcripts are significantly upregulated in both ZH and CabZ mutant GV oocytes (Fig. S2B). Recently, Ishiguro et al., 2017, described Zscan4 expression during the process of oogenesis, and noted that Zscan4 protein shows different staining patterns in NSN or SN types of GV oocytes (diffuse and stronger signals in NSN to spotty and weaker signals in SN) ^18^. Additionally, the authors also observed that Zscan4 appearance correlates with the RNA Polymerase II-mediated transcriptional status during the NSN-to-SN transition. Thus, we hypothesised that the mis-upregulation of *Zscan4* observed in the ZH and CabZ mutant oocytes reflects an aberrant an transcriptional status. To test this hypothesis, we investigated Zscan4 protein level and global transcription activity in ZH and CabZ oocytes.

We performed immunofluorescence (IF) analysis using a newly developed Zscan4 antibody which recognises the truncated forms of Zscan4 including Zscan4a, Zscan4b, and Zscan4e ^18^, and confirmed a significantly higher level of Zscan4 expression in both ZH (142%, compared to ZH Het group set at 100%; p=0.0028; Fig. 3A) and CabZ mutant oocytes (151%, compared to CabZ Het group of 100%; p=0.0143; Fig. 3B). We also noted that *Zscan4* along with other 2-cell-expressed genes (e.g. *Ddit3* and *Gm5039*) persistently showed higher expression in both ZH and CabZ mutant MII stage oocytes, a finding as validated by qRT-PCR (Fig. S3A). This result is in accordance with previously published datasets ^19^ of the Hira conditional mutant MII oocytes driven by both Zp3-Cre and Gdf9-Cre recombination approaches (Fig. S3B).

**Figure 3.**
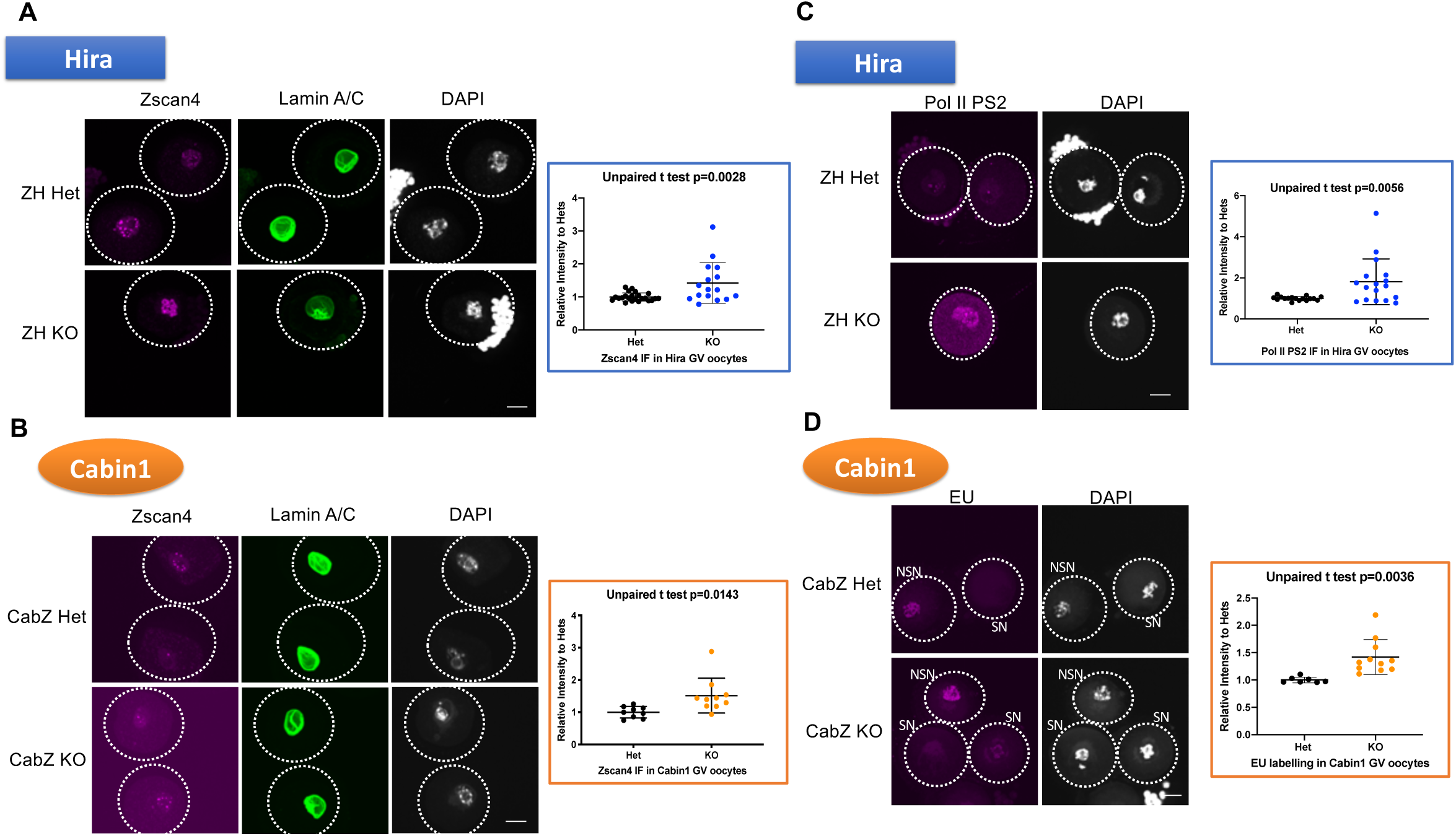
Loss-of-Hira complex disrupts global transcription silencing and de-represses Zscan4 expression and fails in GV stage oocytes. (A) Zscan4 is upregulated in Hira mutant GV oocytes. Immunofluorescent images (left panel) and quantification (right panel) of Zscan4 and Lamin A/C of Hira mutant (ZH KO) and heterozygous control (ZH Het) oocytes. (B) Zscan4 is upregulated in Cabin1 mutant GV oocytes. Immunofluorescent images (left panel) and quantification (right panel) of Zscan4 and Lamin A/C of Cabin1 mutant (CabZ KO) and heterozygous control (CabZ Het) oocytes. (C) Hira mutant oocytes failed to silence global transcription. Immunofluorescent images (left panel) and quantification (right panel) of transcription status monitored by RNA polymerase II phosphorylated Ser2, an active transcription elongation marker. (D) Cabin1 mutant oocytes failed to silence global transcription. Nascent RNA transcription was detected by EU (5-ethynyl uridine) labeling assay. Left panel: EU images; right panel: quantification of EU. SN: non-surrounded nucleolus oocytes; NSN: non-surrounded nucleolus oocytes. Scale bar: 25μm

Next, to monitor the global transcription status in Hira complex mutants, we performed IF of Pol II PS2, a marker of active transcription elongation, on ZH oocytes. Significant upregulation was observed in the ZH KO GV oocytes compared to ZH Het oocytes (180%, compared to ZH Het group of 100%; p=0.0056; Fig. 3C). We then applied 5-Ethynyl-uridine (EU) labelling to assess the nascent RNA synthesis in CabZ oocytes. We showed that the global level of transcription is also elevated in the CabZ mutant oocyte (138%, compared to CabZ Het group of 100%; p=0.0036; Fig. 3D). ZH mutant oocytes also revealed the increasement detected by EU labelling (113%, compared to ZH Het group of 100%; data not shown).

Our work demonstrated upregulated genes from RNA-seq, as well as Zscan4 derepression, and also the aberrant transcription status as shown by IF of Pol II PS2 and EU labelling. This combination of results supports our hypothesis and demonstrates that the Hira complex is essential for the silencing of global transcription, one of the hallmark features of developmental competence in oocytes.

### Two key repressive histone marks, H3K4me3 and H3K9me3 fail to be established in Hira and Cabin1 mutant GV oocytes

It has been concluded that transcription silencing in SN oocytes is highly associated with global chromatin remodelling ^20, 21, 22, 23^, another key feature of a competent oocyte ^24^. Recent research indicates that two main histone repressive mark in this context, H3K9me3 and H3K4me3 ^25, 26^, are responsible for the transcription quiescence in SN oocytes ^23^.

The Hira complex is responsible for H3.3 incorporation and H3.3 has been shown as the only H3 histone subtype incorporated in fully-grown oocytes ^19^. We therefore sought to determine whether the pattern of histone marks is altered in ZH and CabZ mutant oocytes.

IF of H3K4me3 shows decreased levels in ZH KO oocytes (66%, compared to ZH Het group of 100%; p=0.0137; Fig. 4A) as well as in CabZ KO oocytes by IF (54%, compared to CabZ Het group of 100%; p<0.0001). Western blot analysis (Fig. 4B) confirmed these differences. Further analyses of the di-methyl group of H3K4, H3K4me2, also clearly showed lower levels of H3K4me2 in both ZH (57%, compared to ZH Het group of 100%; p=0.0002; Fig. S4A) and CabZ mutant oocytes (64%, compared to ZH Het group of 100%; p=0.0015; Fig. S4B). These data suggest that methylation of H3K4 is under the control of Hira mediated H3.3 incorporation.

**Figure 4.**
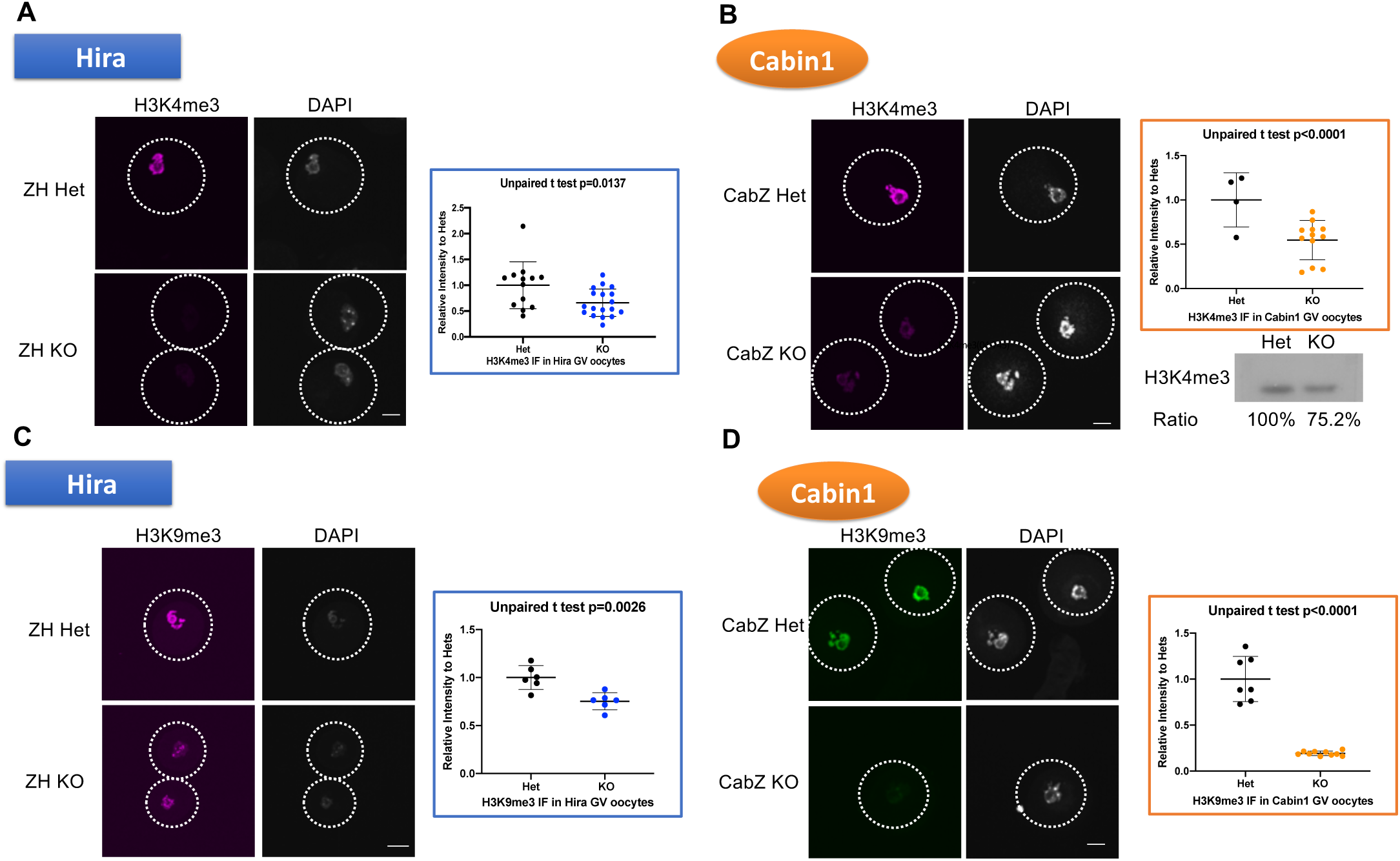
Hira complex is required to establish of repressive histone marks in GV oocytes. (A) Hira mutant oocytes fail to establish H3K4me3 mark. Immunofluorescent images (left panel) and quantification (right panel) of H3K4me3 in Hira mutant (ZH KO) and heterozygous control (ZH Het) oocytes. (B) Cabin1 mutant GV oocytes fail to establish H3K4me3 mark. Immunofluorescent images (left panel), quantification (upper right panel), and western blot (lower right panel) of H3K4me3 in Cabin1 mutant (CabZ KO) and heterozygous control (CabZ Het) oocytes. (C) Hira mutant GV oocytes fail to establish H3K9me3 mark. Immunofluorescent images (left panel) and quantification (right panel) of H3K9me3 in Hira mutant (ZH KO) and heterozygous control (ZH Het) oocytes. (D) Hira mutant GV oocytes fail to establish H3K9me3 mark. Immunofluorescent images (left panel) and quantification (right panel) of H3K9me3 in Cabin1 mutant (CabZ KO) and heterozygous control (CabZ Het) oocytes. Scale bar: 25μm

We also examined another panel of repressive histone modifications, those affecting methylation of H3K9. The level of H3K9me3 was reduced in both ZH KO oocytes (76%, compared to ZH Het group of 100%; p=0.0026; Fig. S4A) and CabZ KO oocytes (19%, compared to CabZ Het group of 100%; p<0.0001; Fig. 4D). The levels of H3K9me2 also revealed a substantial decrease in the ZH KO (73%, compared to ZH Het group of 100%; p=0.0109; Fig. S4C) and CabZ KO (79%, compared to ZH Het group of 100%; p=0.0022; Fig. S4D).

As shown in Fig. S4 E-F, downregulation of the heterochromatin mark (HP1β) was observed in both ZH (57%, compared to CabZ Het group of 100%; p<0.0001; Fig. S4E) and CabZ KO (21%, compared to CabZ Het group of 100%; p<0.0001; Fig. S4F) oocytes. These results strongly indicate that the loss of either Hira or Cabin1 in oocytes leads to reduced levels of repressive histone marks.

Moreover, changes of the panH3 level (75%, compared to CabZ Het group of 100%; p<0.0001 by IF and western blot; Fig. S4G) and H3K27me3 (177%, compared to CabZ Het group of 100%; p<0.0001; Fig. S4H) but not H3K27ac (Fig. S4H) suggest that histone marks may be more broadly altered in the CabZ mutant oocytes (Fig. S4H). This finding is similar to that previously reported in Hira mutants^19^.

### Cabin1 is required for chromatin compaction in GV oocytes

The loss of repressive histone marks in Hira and Cabin1 oocytes prompted us to further investigate possible alterations of chromatin structure. First, we examined the overall chromatin accessibility using a DNaseI-TUNEL assay ^19, 15^ CabZ KO oocytes had significantly stronger TUNEL signals (146%, compared to CabZ Het group of 100%; p=0.0003; Fig.5A) revealing that the overall chromatin architecture had been converted to a less condensed status, echoing previous observations in Hira mutants ^19^. These results also imply that the loss of key repressive histone marks closely reflects the perturbation of chromatin structure.

To further map the chromatin landscape of Hira complex mutant oocytes on a genome-wide scale, we performed Assay for Transposase-Accessible Chromatin using sequencing (ATAC-seq) (Fig. 5A, right panel). In the oocytes, the majority of ATAC-seq peaks were detected in the intergenic and intron regions (Fig. 5A, left panel). ATAC-seq peaks were also preferentially enriched both at the transcription start sites (TSS) and transcription terminatation sites (TTS) (Fig. 5A, right panel), suggesting they might act as promoters and enhancers respectively to regulate gene expression.

**Figure 5.**
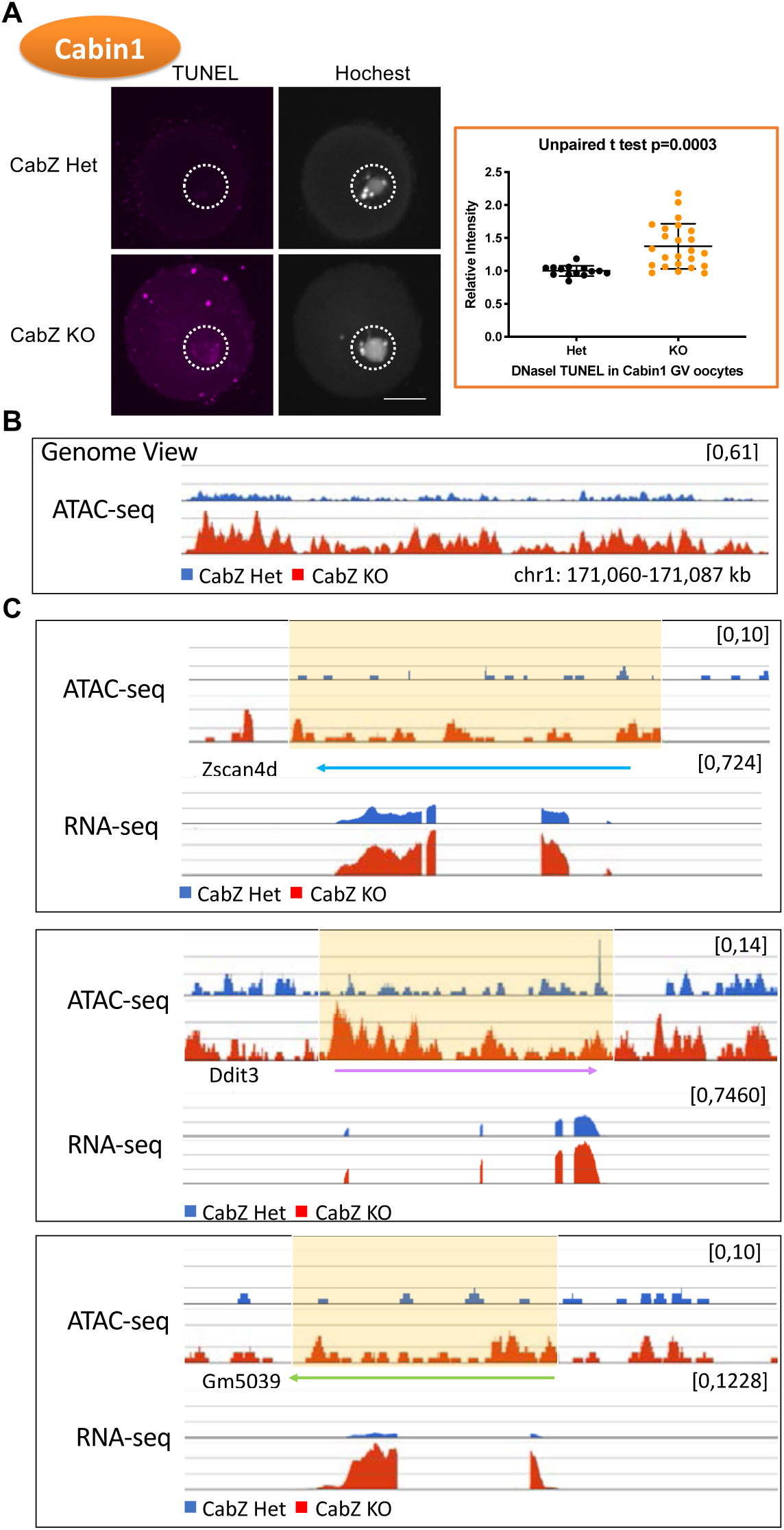
Cabin1 is required to shape the condensed chromatin landscape in GV oocytes. (A) Global chromatin accessibility is increased in Cabin1 mutant oocytes by DNase-TUNEL assay. Left panel: Confocal images of DNase-TUNEL assay; right panel: Quantification of DNase-TUNEL assay. Scale bar: 25μm. (B) Representative genome browser view showing Cabin1 depletion resulted in increasing chromatin openness on the locus of chromosome1 171,060-171,087 Kb. (C) Genome browser views of the combination of RNA-seq and ATAC-seq profiles of Zscan4 and Ddit3, and Gm5039.

There is a sharp contrast between the overall chromatin accessibility of CabZ Het and KO GV oocytes. Specifically, we observed a shift to increasingly accessible chromatin in the CabZ KO oocytes, as shown in the representative genome browser views of detected ATAC-seq peaks in the chr1 171,060-171,087Kb (Fig. 5B) and selected genes, *Zscan4* (Fig. 5C) and *Kera* (Fig. S5B). To further determine whether gene accessibility is associated with corresponding gene expression, we performed integrate analysis using the ATAC-seq and the RNA-seq datasets. We particularly focused on the main candidate genes derepressed in both Hira and Cabin1 mutant oocytes (i.e. *Zscan4c, d, Ddit3*, and *Gm5039*; Fig. 5C and Fig. S5C). We noted distinctly elevated accessible chromatin around the TSS and within the gene bodies of these genes in Cabin1 mutant oocytes (Fig. 5C, Fig. S5C), which coincides with their increased gene expression at the same stage.

In summary, we propose that Cabin1 modulates chromatin architecture by controlling the deposition of key repressive histone signatures and maintenance of a globally condensed chromatin architecture. This is a hallmark feature of developmental competence in GV oocytes. Depletion of Cabin1 causes the loss of repressive histone marks (i.e. H3K4me3 and H3K9me3) coupled with abnormal openness of accessible chromatin loci that leads to misregulation of maternal transcripts (e.g. *Zscan4*).

### Prolonged expression of Zscan4 results in loss of developmental competence and mimics the phenotype of Cabin1 mutant 2-cells

To clarify how histone marks regulate the derepression of Zscan4, we initially performed a series of pharmacological inhibitor experiments. We used a) BIX-01294, an inhibitor of G9a histone lysine methyltransferase ^27^, which decreases the level of H3K9me2; b) MM-102, an inhibitor for MLL1, the core complex of H3K4 histone lysine methyltransferase, which results in decreased levels of H3K4me3 ^28^; and c) the combined treatment of BIX-01294 with MM-102 (Fig. S6A). We cultured wild-type GV oocytes with these inhibitors to monitor whether the level of Zscan4 increases after withdrawal of the repressive histone marks. We firstly confirmed the efficiency of the inhibitors. In treatment a), the level of H3K9me2 was reduced in the BIX-01294 treated group (29%, compared to DMSO group of 100%; p<0.0001); in treatment b), the level of H3K4me3 was reduced in the MM0102 treated group (71%, compared to DMSO group of 100%; p=0.0237); and in treatment c), the combined treatment successfully induced reduction of the level of H3K9me2 (51%, compared to DMSO group of 100%; p<0.0001); and also reduced the level of H3K4me3 (73%, compared to DMSO group of 100%; p=0.0013; Fig. S6A). Importantly, we observed elevated expression of Zscan4 by qRT-PCR in all three treatments (Fig. S6A). These results suggest that the repressive histone marks (i.e. H3K4me3 and H3K9me3) act as upstream regulators to modulate the proper expression of Zscan4.

To further determine whether mis-regulation of Zscan4 in the oocyte directly contributes to consequent developmental arrest, we forced the expression of Zscan4 in wild-type GV oocytes.

We used the same setup for Zscan4d RNA injection in GV oocytes as for our MO injections followed by *in vitro* maturation (IVM), selection of MII oocytes for parthenogenetic activation, and subsequent *in vitro* culture (Fig. 6A). Successful overexpression was validated by qRT-PCR and IF of the tagged Flag Zscan4 protein (Fig. S6B-C). The developmental potential of the embryos was determined by the progression to morula-to-blastocyst stage embryos. There was no significant difference in the proportion of 2-cell cleavage rate between the Zscan4 overexpression group and the control group (GFP injected). By contrast, the developmental rate to morula-to-blastocysts was significantly reduced in the Zscan4 overexpression group (31%, compared to control GFP group of 53%; p=0.0261; Fig. 5B). However, very little changes of key histone marks (H3.3, H3k4me3, and H3K9me3) was observed by IF in the Zscan4 overexpressed oocytes (Fig.6D). This suggests that the alterations of key repressive histones in both ZH and CabZ mutants is not triggered by the upregulation of Zscan4.The mechanism of Zscan4 expression is more likely mediated by upstream histone modifications pathways. These results provide evidence that timely expression of Zscan4 is critical for embryogenesis.

**Figure 6.**
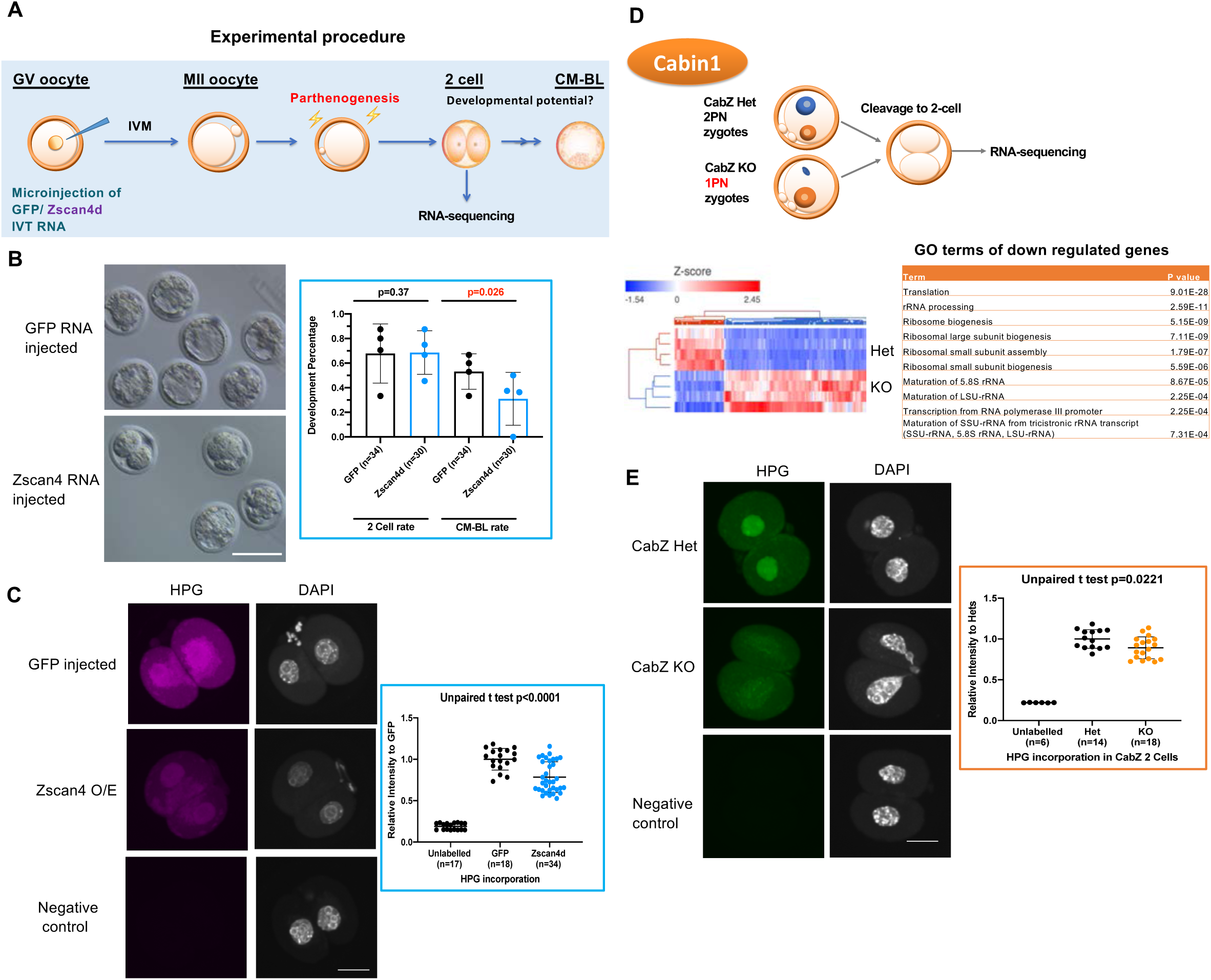
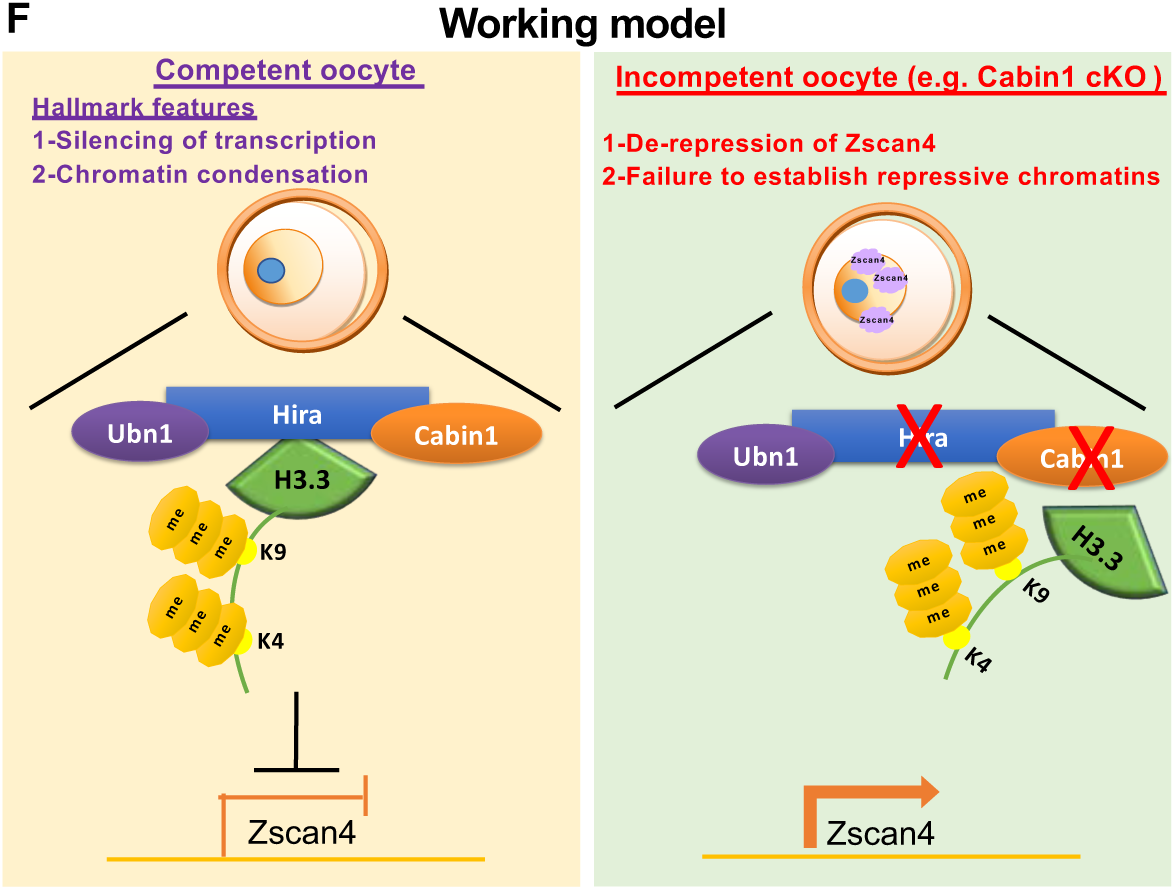
Overexpression of Zscan4 impairs developmental incompetence. (A) Experimental procedure of over-expression of Zscan4 in oocytes and developmental/molecular analyses. RNA injected oocytes were matured *in vitro*, parthenogenetically activated by SrCl_2_ for 5 hours and cultured *in vitro*. 2-cell formation and blastocyst development potential were monitored. Parthenogenetically activated 2-cells of Zscan4 and GFP RNA injected oocytes were sent for RNA-sequencing analysis. (B) Zscan4d over-expressed oocytes compromised developmental potential after parthenogenesis and *in vitro* culture. Left panel: Developmental progression of GFP injected and Zscan4 injected embryos at embryonic day 4 (E4). Right panel: quantification of 2-cell cleavage and morula-to-blastocyst formation rate. Scale bar: 100μm. (C) Global protein synthesis is repressed in Zscan4d over-expressed embryos by HPG incorporation assay. Left panel: Images of HPG incorporation in Zscan4 overexpression and controls (GFP injected and unlabeled) embryos. Right panel: Quantification of HPG incorporation assay. Scale bar: 25μm. (D) Gene Ontology of RNA-seq result of Cabin1 mutant 2-cells showed that regulation of translation related terms are among the top downregulated lists. (E) Global protein synthesis is repressed in the Cabin1 mutant 2-cell embryos by HPG incorporation assay. Left panel: Images of HPG incorporation in Cabin1 mutant and controls (Cabin1 Heterozygous and unlabeled) embryos. Right panel: Quantification of HPG incorporation assay. Scale bar: 25μm. (F) A working model describes the mechanism controlling oocyte competence by Hira mediated epigenome and transcriptome remodeling.

Knockdown/knockout of maternal genes that are critical for zygotic genome activation and early embryogenesis (e.g. Brg1, Mll2, and Kdm1a) often induces an overall reduction of initiation of translation ^11, 29^. Indeed, 2-cell stage parthenogenetic embryos derived from overexpressed Zscan4 oocytes displayed a significant reduction of global protein synthesis compared to 2-cell stage embryos from GFP RNA injected oocytes as assessed by the homopropargylglycine (HPG) incorporation assay (73%, compared to control GFP group of 100%; p<0.0001; Fig. 6C).

Intriguingly, comparison between RNA-seq data from the CabZ KO and CabZ Het 2-cells revealed that top-ranked GO terms of DE down-regulated genes appeared to be classified as translation including rRNA processing, ribosome assembly and ribosome biogenesis (Fig. 6D). HPG incorporation assays also showed a significantly lower protein production in the CabZ KO compared to CabZ Hets 2-cells (89%, compared to CabZ Het group of 100%; p<0.0221; Fig. 6C), which agrees with the Zscan4 phenotype. The consistency of impairment on protein production of CabZ mutants and Zscan4 overexpression 2-cell embryos indicated that there is a mutual deficit of the initiation of ZGA.

Moreover, RNA-sequencing results of Zscan4 overexpressed 2-cells revealed that transcripts involved in “Regulation of transcription”, “Blastocyst development” and “Ribosome small subunit assembly” were overrepresented among the downregulated GO terms (Fig. S6E). Above all, Rps19, an essential ribosomal protein that is crucial for blastocyst development ^30^ was shared in downregulated genes of CabZ and Zscan4 overexpressed 2-cells. qRT-PCR also confirmed the downregulation of *Rps19* confirming the results of RNA-seq (Fig. S6F).

Therefore, likely failure to begin ZGA associated with the decline of overall protein production of the CabZ KO 2-cells is likely due to the derepression of Zscan4 in oocytes. This in turn hampers embryonic development.

Taken together, our results can be accommodated in a model (Fig. 6F) whereby competent oocytes have two hallmark features of 1) silence of transcription and 2) regulation of overall chromatin condensation by Hira complex mediated H3.3 incorporation. This state is associated with repressive marks of H3K4me3 and H3K9me3 that suppress activation of a subset of early embryonic genes (e.g. Zscan4). In contrast, incompetent oocytes, such as those resulting from maternal disruption of Hira or Cabin1, fail to establish marks of H3.3 incorporation with K4me3 and K9me3 modifications. As a consequence, an aberrant chromatin landscape results in premature activation of gene expression (e.g. derepression of Zscan4 and other embryonic genes) eventually leading to a developmental arrest.

## Discussion

Using several complementary approaches, our data demonstrate that the Hira-complex plays an essential role in the development of the oocyte competence to sustain embryo devleopment at the oocyte-to-embryo transition. Mechanistically, this complex is involved in shaping the chromatin by promoting incorporation of H3.3 enriched with H3K4me3 and H3K9me3 marks. The methylated H3.3 reduces chromatin accessibility and represses transcription of down-stream early embryonic genes such as *Zscan4, Ddit3* and *Gm5039*. Failure to repress these genes causes delayed effects after fertilisation/parthenogenesis in the embryo. Consistent with this conclusion, overexpression of Zscan4 prevents the proper initiation of zygotic genome activation and causes a developmental arrest that recapitulates the key phenotypes of loss-of-function of Hira-complex mutants. Thus, enforcing a repression of transcription in quiescent GV oocytes is an indispensable event that prepares for the programmed changes in gene expression in the zygote that are critical for establishing totipotency.

Despite identification years ago of two distinctive features of competent oocytes, chromosomal condensation and formation of heterochromatin rims ^22, 31^ as well as transcriptional silencing ^32^, the underlying regulatory mechanisms repressing transcription have remained elusive. With our study, we demonstrate that both Hira and Cabin1 mutant oocytes lack H3K4me3/HK9me3 marks. Therefore, the Hira-complex plays an essential role in remodelling of the chromatin at this critical transition of oocyte development. Altered H3.3 methylation that follows disruption of the Hira complex is associated with increased chromatin accessibility assessed by ATAC-seq and altered transcription measured by RNA-seq, with most of affected genes being upregulated. These observations strongly argue that the repressive H3.3 methylation marks are indispensable to induce the transcription silencing observed at this stage.

Failure of incorporation of methylated H3.3 in the oocyte leads to an early embryo arrest indicating a loss of competence to develop as embryos. Our data challenge the idea proposed by Zuccotti et al.^22^ that the heterochromatin rim can be used as the unequivocal selection marker for competent oocytes. Both Hira and Cabin1 mutant oocytes have lost their developmental competence despite maintaining the heterochromatin rim structure. It is likely that, in addition to forming the heterochromatin rim structure, additional mechanisms in the oocyte are necessary to reach a higher degree of chromatin condensation to complete the final stage of the NSN-SN transition. According to this view, oocyte repressive histone modifications, including H3K4me3 and H3K9me3, may serve as the “competent oocyte” histone code deposited by the Hira complex. Genome-wide profiling by Hi-C ^33^ and/or in combination with higher resolution imaging technique such as electron spectroscopic imaging (ESI) ^34^ should add further support to this idea.

Our data provide insight into the molecular mechanism involved in this generalised repression of transcription enforced in the oocyte prior to meiotic re-entry. Cabin1 has been identified as a vertebrate-specific subunit of Hira complex (there is no Cabin1 homolog in fly). Here, we identified Cabin1 as a new maternal factor that is critical for fecundity. Cabin1 homozygous null mutants exhibiting embryonic lethality during organogenesis has been reported ^35^. Cabin1 roles not only include binding to Calcineurin and myocyte enhancer factor 2 (MEF2) in T cell development, but also chaperoning H3.3 incorporation via binding to trimerised Hira ^8^, the embryonic lethality phenotype in Cabin1 mutants likely results from insufficient incorporation of H3.3. Morevoer, it has been shown that Cabin1 recruits Suv39H1, a major H3-K9 methyltransferase required to repress Myocyte enhancer factor 2 transcription activity in T cells ^36^. In agreement with this view, we also observed a decreased level of H3K9me2/3 and HP1 in Cabin1 mutant oocytes. This finding supports the notion that Cabin1 likely has a similar mechanism of action in oocytes to suppress the developmental genes programmed to be expressed at later stages. Similarly, Cabin1 may play a function in the PRC1/PRC2 or Suv39h1/Suv39h2 mediated maternal-to-zygotic transition ^37, 38^.

Among the GO terms of the differentially up-regulated genes in Hira mutant oocytes, the candidate Zscan4 was selected in the term “Telomere maintenance via telomere lengthening”. However, we surmised that Zscan4 might not specifically function on telomeres because oocytes have shorter telomeres than somatic cells, and their telomeres increase in length later during early cleavage development ^21^. Therefore, we focused on other functions for Zscan4 during the oocyte-to-embryo transition. Zscan4 upregulated status in the Hira and Cabin1 mutant oocytes reflected the deficit in transcriptional repression.

Zscan4 plays pivotal roles during development ^39^, cell fate fidelity ^16,^, regulation of transcription ^40, 41^, telomere and genomic stability ^42^ and is a specific marker of 2-cell like embryonic stem cells ^42, 43^. Also, misregulation of Zscan4 accompanies failure of progression through embryo development ^15, 44, 45^, and imbalance of protein synthesis ^40^. In agreement with these findings, overexpression of Zscan4 compromises the oocyte developmental potential and mimics the key phenotypes of Cabin1/Hira mutants. In both cases, advanced developmental stages of the embryo are affected owing to the impairment of ZGA initiation and protein production. It will be of interest to further identify Zscan4’s downstream targets.

Intriguingly, embryonic Zscan4 expression is under tight control by chromatin regulators such as PRC1 complex (i.e. Ring1/Rnf2; ^46^) and methylation of H3K9 ^44^. Ring1/Rnf2-deficiency in oocytes leads to the precocious expression of Zscan4 and of lineage markers such as *Sox2, Klf4*, and *Eomes* which are thought to be expressed in later embryonic stages ^46^. We found that the BIX-01294 and MM-102 inhibitors also induced upregulation of *Zscan4* in oocytes, suggesting that *Zscan4* expression is downstream of the regulation of H3K4 and H3K9 by methylation. On the other hand, overexpression of Zscan4 did not alter histone marks, supporting the notion that it acts downstream of these histone modifications. All these findings taken together provide a strong link between the H3.3, the Hira complex and the mechansims of gene silencing occuring at this stage of oocyte development.

In summary, we demostrate that the H3.3 chaperone Hira complex is essential for oocyte developmental competence by shaping its epigenome and transcriptome. The oocyte withholds totipotent/pluripotent potential by repressing a subset of early embryonic genes via deposition of repressive histones. Even though the chromatin condensation process is conserved in most mammalian oocytes, diverse dynamics have been noticed. For example, 4 stages of chromatin condensation steps had been observed in bovine oocytes ^47^. Similar properties shared by bovine and human oocytes would make it worthwhile to re-examine the levels of H3K4me3, H3K9me3, and also Zscan4 in these species. Also, it will be of interest in the future to explore whether Cabin1, in cooperation with Hira, is involved in male pronucleus formation and nucleosome assembly in both mouse and human (manuscript in preparation). Moreover, further research will be critical for the identification of Zscan4 targets on the regulation of chromatin condensation and the development of epigenetic biomarkers for competent oocytes, both of which may provide new routes to improving assisted reproductive technologies.

## Supplement Figure Legends

Figure S1. Experimental approaches for investigation of the maternal role of the Hira complex.

(A) Ubn1 knockdown and developmental assay. GV oocytes were microinjected with Ubn1 and control antisense morpholino oligos. Injected oocytes were matured *in vitro*, parthenoactivated by SrCl2 for 5 hours and cultured *in vitro*. 2-cell formation and blastocyst development potential were monitored.

(B) Strategy of generation of knockout Hira in the oocytes. Exon 6 and 7 regions (this study; ^48^) of Hira were deleted in the oocyte using Zp3-cre recombination approach. Previous paper published by us ^10^ where we deleted the exon 4 region.

(C) Strategy of generation of knockout Cabin1 in the oocytes. Exon 4 region of Cabin1 was deleted in the oocyte using Zp3-cre recombination approach.

(D) Knocking down of Ubn1 in the oocyte impaired oocyte maturation potential. MII oocyte maturation rate was monitored in Ubn1 knockdown oocytes.

(E) Knocking down of Ubn1 in the oocyte impaired embryo cleavage. 2-cell formation rate was monitored in Ubn1 knockdown oocytes.

(F) Maternal H3.3 is required for preimplantation development of parthenogenetically activated embryos. Upper panel: H3.3 knockdown approach was similar as Ubn1 knockdown. Lower panel: 2-cell formation and morula-to-blastocyst rate of H3.3 knockdown embryos was impaired. Scale bar: 100μm.

Figure S2. Bioinformatic analyses of RNA-seq results of Hira and Cabin1 mutant oocytes.

(A) Minimal overlapping genes of either Hira or Cabin1 differentially expressed genes with growing oocytes (surrounded-nucleus, SN).

(B) A subset of 2 cell specific genes derepressed in both Hira and Cabin1 mutant GV oocytes. Left panels: Venn-diagram showed the overlapping genes which are upregulated both in the Hira and Cabin1 mutant GV oocytes. Right panel: qRT-PCR confirmed the selective upregulated genes, Zscan4d, Gm5039, and Ddit3, in both Hira and Cabin1 mutant GV oocytes.

Figure S3. Zscan4 is upregulated in the Hira and Cabin1 MII oocytes.

(A) Zscan4 is upregulated in both Hira and Cabin1 mutant MII oocytes. qRT-PCR of Zscan4, Ddit3, and Gm5039 expression in Hira and Cabin1 mutant MII oocytes. Expression was normalised to heterozygous controls.

B) List of Zscan4 fold changes of Hira mutant MII oocytes from previously published dataset (Nashun et al.,).

Figure S4. Hira complex is required for establishment of repressive histone and heterochromatin marks in GV oocytes.

(A) Hira mutant GV oocytes fail to establish H3K4me2 mark. Immunofluorescent images (left panel) and quantification (right panel) of H3K4me2 in Hira mutant (ZH KO) and heterozygous control (ZH Het) oocytes.

(B) Cabin1 mutant GV oocytes fail to establish H3K4me2 mark. Immunofluorescent images (left panel) and quantification (right panel) of H3K4me2 in Cabin1 mutant (CabZ KO) and heterozygous control (CabZ Het) oocytes.

(C) Hira mutant GV oocytes fail to establish H3K9me2 mark. Immunofluorescent images (left panel) and quantification (right panel) of H3K9me2 in Hira mutant (ZH KO) and heterozygous control (ZH Het) oocytes.

(D) Cabin1 mutant GV oocytes fail to establish H3K9me2 mark. Immunofluorescent images (left panel) and quantification (right panel) of H3K9me2 in Cabin1 mutant (CabZ KO) and heterozygous control (CabZ Het) oocytes.

(E) Hira mutant GV oocytes failed to establish HP1 mark. Immunofluorescent images (left panel) and quantification (right panel) of HP1 in Hira mutant (ZH KO) and heterozygous control (ZH Het) oocytes.

(F) Cabin1 mutant GV oocytes failed to establish HP1 mark. Immunofluorescent images (left panel) and quantification (right panel) of HP1 in Cabin1 mutant (CabZ KO) and heterozygous control (CabZ Het) oocytes.

(G) Overall histone H3 level was decreased in Cabin1 mutant oocytes. Immunofluorescent images (left panel), quantification (upper right panel), and western blot (lower right panel) of H3K4me3 in Cabin1 mutant (CabZ KO) and heterozygous control (CabZ Het) oocytes.

(H) The level of H3K27me3 mark was upregulated in Cabin1 mutant GV oocytes while H3K27ac mark remained. Immunofluorescent images (left panel) of H3K27me3 and quantification (right panel) of H3K27me3 and H3K27ac in Cabin1 mutant (CabZ KO) and heterozygous control (CabZ Het) oocytes. Scale bar: 25μm

Figure S5. Global chromatin accessibility is increased in Cabin 1mutant GV oocytes

(A) Principal Component Assay of ATAC-seq of Cabin1 mutant GV oocytes.

(B) Genome browser views of the ATAC-seq profiles of a selective gene, Kera.

(C) Genome browser views of the combination of RNA-seq and ATAC-seq profiles of Zscan4c.

Figure S6. Zscan4 expression is regulated by repressive histone marks of H3K4me3 and H3K9me3.

(A) Histone methyltransferase inhibitor experiments confirmed that Zscan4 expression is modulated by H3K4me3 and H3K9me2 modifications. Oocytes were treated with BIX-01294, MM-102, and combined for 48hrs. qRT-PCR was used to confirm the upregulation of Zscan4 after inhibitor treatment. IF was used to confirm the inhibition of target histone marks.

(B) qRT-PCR validation of Zscan4 overexpression in the oocytes. RNA concentration of low Zscan4 (300ng/μl) and high (1,500ng/μl).

(C) Zscan4 overexpression was confirmed by immunofluorescence using antibody against Flag-tag. Scale bar= 25μm.

(D) Over-expression of Zscan4 was unable to alter the level of H3.3, H3K4me3 and H3K9me3.

(E) RNA-seq result of overexpressed Zscan4 oocytes at 2-Cell stages. Left panel: Gene Ontology of RNA-seq result of overexpressed Zscan4 2-cells showed that regulation of transcription, blastocyst development, and ribosomal assembly related terms are among the top downregulated lists. Right panel: Venn-diagram represented the co-downregulated differentially expressed genes of Cabin1 mutant and overexpressed Zscan4 oocytes at 2-Cell stages.

(F) qRT-PCR of CabZ KO and Zscan4 overexpressed 2-cells confirmed the downregulation of Rps19.

## Material and Methods

### Animals

Mouse experiments were approved by the University of Edinburgh’s Animal Welfare and Ethical Review Board (AWERB) and carried out under the authority of a UK Home Office Project License. C57BL/6 wild type mice were purchased from Charles River Laboratories. The mouse lines used in this study carried conditional floxed alleles for Cabin1and for Hira on C57BL/6 backgrounds (both were provided from Prof Peter Adams previously in Beatson Institute, UK, now relocated to Sanford Burnham Prebvs, USA.). Floxed Cabin1 (Cabin1^fl/fl^) and floxed Hira (Hira^fl/fl^) mouse lines were crossed with a Zp3-cre mouse line (deVries *et al*, 2000; provided from Prof Petra Hajkova, Imperial College of London) that expresses cre recombinase in the female germline to generate heterozygous and homozygous mutant oocytes.

### Oocyte/embryo culture and micromanipulation

Female mice were superovulated by administration of 7.5 I.U. pregnant mare serum gonadotrophin (PMSG from Prospec Protein Services, USA) and 48 hours later fully-grown oocytes were isolated. Oocytes were cultured in M16 medium at 37°C, 6%CO_2_ and 5%O_2_. Parthenogenetically activation was carried out as previous described (Lin et al., 2014), matured MII oocytes were cultured in calcium free CZB medium with 10mM SrCl_2_ for 5 hours.

For zygote collection, 48 hours after PMSG administration 7.5 I.U. human chorionic gonadotrophin (hCG, Chorulon® from Intervet) was injected and the mice mated with C57BL/6 males. The following day zygotes were collected from the oviducts. Embryos were cultured in KSOM medium at 37°C, 6%CO_2_ and 5%O_2_.

Micromanipulation was performed as described previously ^5, 10^. Micromanipulation platform was equipped with a microinjector (FemtoJet 4i, Eppendorf), an inverted microscope (Leica, DMi8), and micromanipulators (Narishige). For Ubn1 knockdown experiments, oocytes were microinjected with control or Ubn1 antisense morpholino oligos (Gene Tools). For Zscan4 overexpression experiments, Zscan4 (Addgene 61831) and GFP (gift from M Anger, IAPG, CAS) *in vitro transcription* RNA were carried out using mMessage mMachine® kit (Life Technologies) and a poly(A) tail was added (Applied Biosystems AM1350). Unincorporated nucleotides were then removed with a RNA clean up kit (Zymo R1015).

### Immunofluorescence

Immunofluorescence (IF) experiments were performed as previously reported (Lin et al., 2013 2014). Images of stained embryos were acquired by a spinning disk confocal (CSU-W1, Yokogawa) on an upright microscope frame (BX-63, Olympus) using a 30 or 60x silicon oil immersion objective (UPLSAPO 60XS2, Olympus). IF signal intensity was quantified using Fiji software. Antibodies used listed in Table S1.

### Western blotting

Oocytes were washed in PBS and frozen to −80°C. Thirty oocytes were lysed in Reducing SDS loading buffer and denatured for 5 min. Proteins were separated by gradient precast 4–12% SDS–PAGE gel (Thermo Fisher) and transferred to Immobilon P membrane (Millipore) by the semidry blotting system. Membranes were blocked by 5% skimmed milk for 1 h and incubated with the following primary antibodies (H3K4me3, H3, and Gapdh) diluted in 1% milk*/*TTBS overnight. The membranes were incubated in secondary antibodies diluted 1:10 000 (Peroxidase Donkey Anti-Rabbit) for 1 h at room temperature. Proteins were visualised by chemiluminescence using ECL (Amersham). Negatives were scanned using a GS-800 calibrated densitometer (Bio-Rad).

### Inhibitor experiments

GV oocytes from wild-type mice were cultured in M16 medium containing IBMX (both Sigma) at 37°C, 6% CO_2_, 5% O_2,_ and supplemented with BIX-10294 (Stratech) at final concentrations of 0.1µM and 1µM ^49^; MM-102 (Cayman Chemicals) at final concentrations of 6µM or 60µM, within the range previously reported ^50^; or a combination of 1µM BIX-01294 and 60µM MM-102.

The inhibitors were dissolved in DMSO (Sigma) and an equivalent concentration was used for the control groups. After 48 hours the surviving GV oocytes were either fixed in 4% paraformaldehyde (Thermo Scientific) for investigation by IF, or frozen at −80°C for RNA isolation and subsequent qRT-PCR.

### Nascent transcription and translation assays

For nascent transcription assays, oocytes were incubated in M16 medium with IBMX supplement with 1mM EU (5-ethynyl uridine) for 2 hrs, fixed in methanol, then a Click-iT RNA Imaging kit (Thermo Fisher Scientific) was utilised and visualisation was by confocal microscopy.

Changes in protein expression were monitored using Click-iT Protein Synthesis Assay kit (Thermo Fisher Scientific). Oocytes and 2 cell embryos were cultured in M16 medium with the methionine analogue HPG at a dilution of 1:500. Newly synthesised proteins incorporating the HPG were visualised by confocal microscopy.

### RNA isolation and qRT-PCR

RNA was extracted from oocytes/embryos using PicoPure RNA isolation kit (Arcturus) and cDNA made using iScript(tm) cDNA Synthesis kit (BioRad). Brilliant III SYBR® green QPCR Master Mix (Agilent Technologies) was used for qRT-PCR and reactions run on a Roche LightCycler®96 real time PCR system. Gene of interest mRNA levels were calculated by comparison with Hprt1 and H2A mRNA levels using REST software. Primers used listed in Tabls S2.

### RNA-seq library preparation and data processing

Ten oocytes collected from each experimental treatments or WT were pooled and cDNA was amplified using Smart-seq2 v4 kit with minor modification from manufacturer’s instructions. Briefly, oocytes were lysed, and mRNA was captured and amplified with the Smart-seq2 v4 kit (Clontech). After AMPure XP beads purification, amplified RNAs were quality checked by using Agilent High Sensitivity D5000 kit (Agilent Technologies). RNA-seq libraries were constructed using Nextera XT DNA Library Preparation Kit (Illumina) and multiplexed using Nextera XT Indexes kit (Illumina). Libraries were quantified by Qubit (Life Technologies) and Tapestation 4200 (Agilent). Pooled indexed libraries were then sequenced on the Illumina HiSeq X platform with 150-bp pair-end reads.

Multiplexed sequencing reads that passed filters were trimmed to remove low-quality reads and adaptors by TrimGalore-0.4.3. Clean reads were aligned to the mouse genome (GRCm38/mm10) by using STAR with default parameters. Approximately 30 million reads per individual cell were generated. Individual mapped reads were adjusted to provide FPKM (fragments per kilobase of exon model per million mapped fragments) values with RefSeq genes as reference. Differential gene expression analysis was performed by a Partek Flow GSA algorithm with default parameters. The genes were deemed differentially expressed if they provided a false discovery rate (FDR) adjusted p-value of <0.05 and fold change >2. DAVID (https://david.ncifcrf.gov) and IPA (Ingenuity Pathway Analysis) were used to reveal the Gene Ontology (GO) and pathways, respectively.

### ATAC-seq library preparation and data analysis

The ATAC-seq libraries of oocytes were prepared as previously described with minor modifications ^14, 51^. Briefly, pooled oocytes were lysed in ice-cold lysis buffer (10mM Tris-HCl (pH 7.4), 10mM NaCl, 3mM MgCl_2_ and NP-40 (0.5%) for 15 minutes on ice to prepare the nuclei. Immediately after lysis, Nuclei were then incubated with the Tn5 transposase (TDE1) and tagmentation buffer at 37°C for 30 minutes with shaking on a thermomixer at 500g. After the tagmentation, 0.5 ul of 10% SDS was added directly into the reaction to end the tagmentation and tagmentated DNA was purified using MinElute Reaction Cleanup Kit (Qiagen). PCR was performed to amplify the ATAC-seq libraries using Illumina TrueSeq primers and multiplex by indexes primers with the following PCR conditions: 72°C for 3min; 98°C for 30s; and 15 thermocycling at 98°C for 15s, 60°C for 30s and 72°C for 3min; following by 72°C 5 min. After the PCR reaction, libraries were purified with the 1.1X AMPure beads (Beckman). Libraries were quantified by Qubit (Life Technologies) and Tapestation 4200 (Agilent). Pooled indexed libraries were then sequenced on the Illumina HiSeq X platform with 150-bp pair-end reads.

All quality assessed ATAC-seq reads were aligned to the mouse reference genome using Bowtie 2.3 with following options: --very-sensitive -X 2000 --no-mixed --no-discordant. Alignments resulted from PCR duplicates or locating in mitochondria were excluded. Only unique alignments within each sample were retained in subsequent analysis. Principle component analysis (PCA) was performed on pairwise correlation data in R with functions prcomp. ATAC-seq peaks were called separately for each sample by MACS2 ^52^ with following options: --keep-dup all – nolambda --nomodel. Peaks in individual samples from the same development stage were subsequently merged using bedtools (https://bedtools.readthedocs.io/en/latest/). For downstream analysis, we normalized the read counts by computing the number of reads per kilobase of bin per million of reads (RPKM). To visualize the ATAC-seq signal in the UCSC genome browser, we extended each read by 250bp and counted the coverage for each base.

### DNaseI-TUNEL assay

Oocytes were permeabilized by 0.5% Triton X-100 in pre-extraction buffer (300mM sucrose, 25mM HEPES, 1M CaCl_2_, 50mM NaCl, 3mM MgCl_2_) for 5 min before digesting with 0.2U/ml of DNase I (NEB). Embryos were then fixed in 4% PFA. TUNEL Assays (Click-iT TUNEL Imaging assay, Thermo Fisher Scientific) were followed according to manufacturer’s instructions. Images were visualised by confocal microscopy.

### Statistical analyses

For analyses of percentage of oocyte maturation, embryonic development, the χ^2^ test was used as previously described (Lin et al., 2013 and 2014). For analyses of IF intensity, signals of EU, DNaseI-TUNEL, and HPG incorporation, two-tailed *t*-test with unequal variance was used. All error bars indicate s.d.

### Data availability

The raw FASTQ files and the data sets generated and analysed during the current study are available in the Gene Expression Omnibus (GEO) (https://www.ncbi.nlm.nih.gov/geo/) under the accession number GSE146894.

## Acknowledgements

This work was supported by MRC Centre Grant MR/N022556/1, the Wellcome Trust-University of Edinburgh Institutional Strategic Support Fund, and grants from Deanery of Clinical Sciences, College of Medicine & Veterinary Medicine of University of Edinburgh to C.-J. L. C.-J. L. is a Royal Society of Edinburgh Personal Research Fellow funded by the Scottish Government. Z.J was supported by the USDA-NIFA grant (2019-67016-29863), the Audubon Center for Research of Endangered Species, and UDSA-NIFA W4171.

We thank P. Hajkova for Zp3-Cre mice, P. Adams for Hira and Cabin1 mice and Ubn1 antibody, M. Ko for Zscan4 antibody also the plasmids of Zscan4 (Addgene 61831) and GFP (Martin Anger, ^53^). We also thank Jaroslava Supolikova and Marketa Hancova in IAPG, CAS for the technical assistance for western blot assay. We thanks the constructive feedbacks from Prof Adele Marston and Prof Miguel Ramalho-Santos.

## Author contribution

C.-J. L.: Conceived the project and designed the experiments performed all embryo manipulation and part of the mouse work.

R. S.: Mouse colony management and performed oocyte experiments.

Z. J. and H. M.: Performed genome-wide experiments and bioinformatic analyses.

A. S.: Performed western blot of histone marks of mouse oocytes.

J. T.: Characterisation of Cabin1 mice.

C.-J. L., Z. J., M.C., and A. S. interpreted the data and assemble the results.

R. S., Z. J., M.C., and C.-J. L. wrote the manuscript with inputs from all authors.

## Notes

### Competing Interest Statement

The authors have declared no competing interest.

