## Supplemental Figures for "The H3.3 chaperone Hira complex orchestrates oocyte developmental competence"

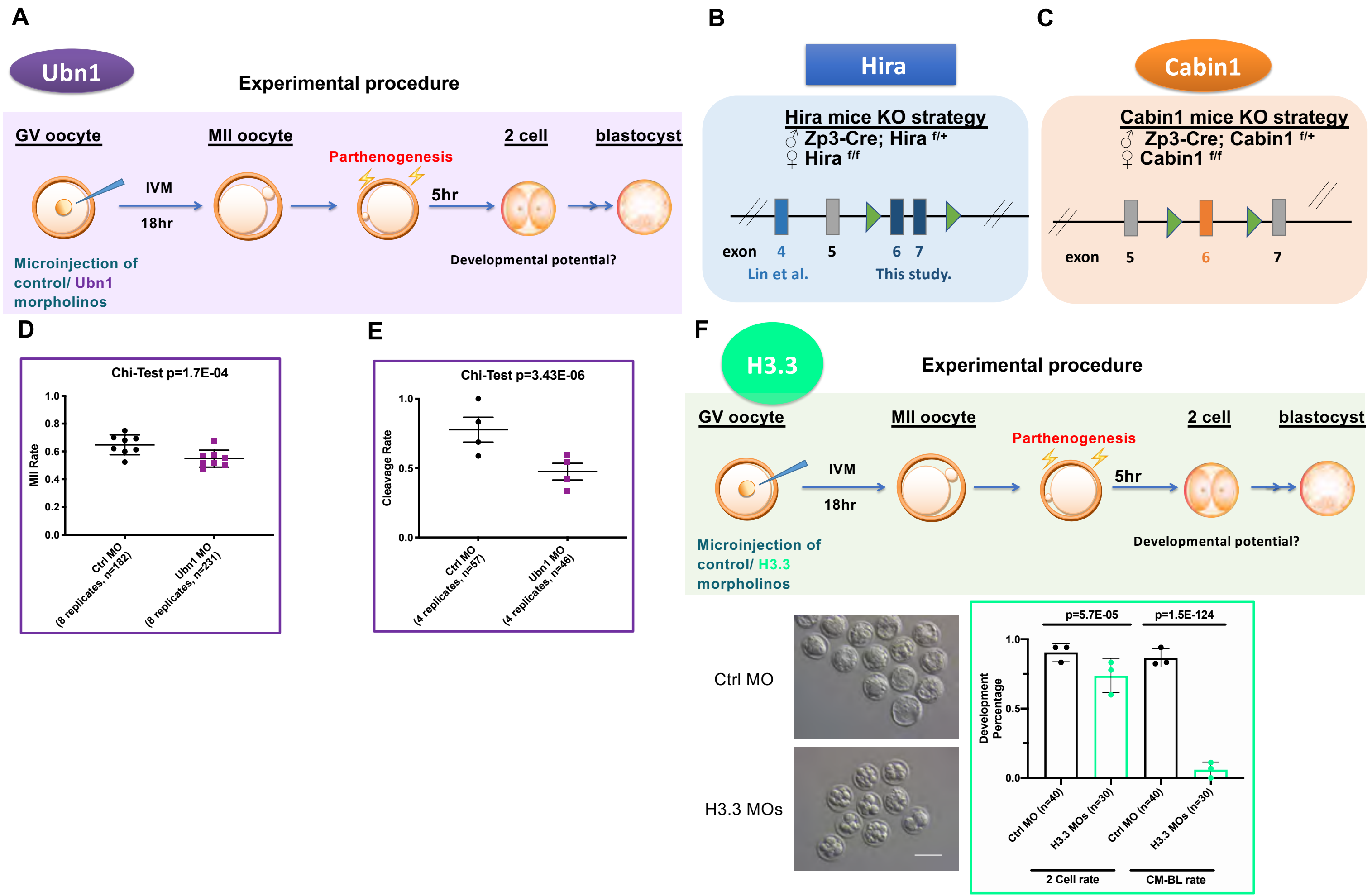


Figure S1 Experimental approaches of investigating the role of the Hira complex in the oocyte.

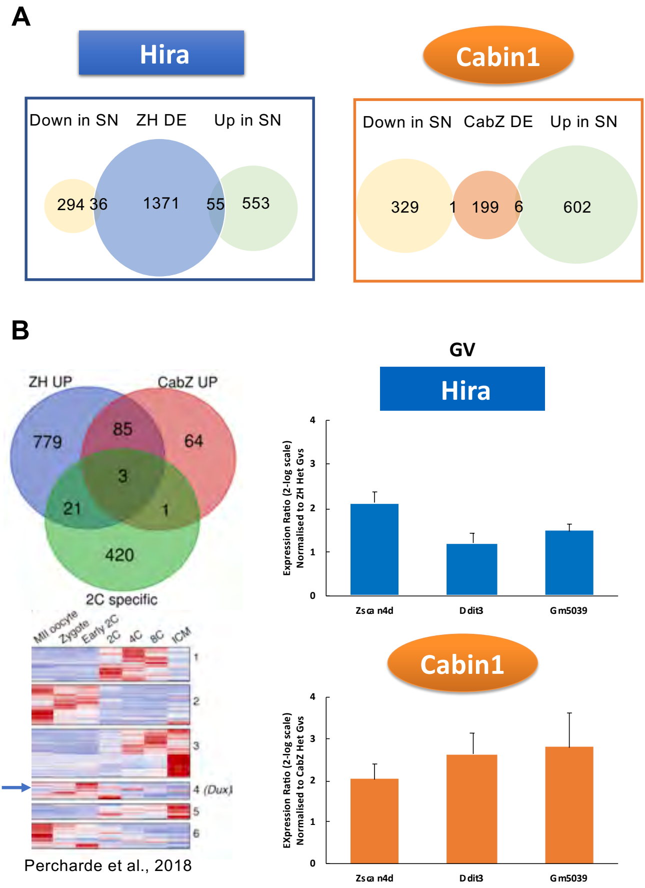


Figure S2 Loss-of-Hira complex in the GV oocytes induce a subset of 2cell specific genes

**A**

**MII**

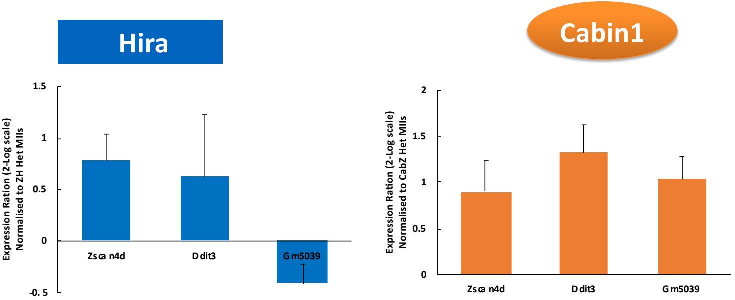


**B**

| logFC | Gdf9-Cre Hira KO | Zp3-Cre Hira KO |
|---|---|---|
| Zscan4c | 4.97 | 1.38 |
| Zscan4d | 4.43 | 2.18 |
| Zscan4f | 4 | 2.29 |
| Ddit3 | 2.39 | 0.465 |
| Gm5039 | NA | NA |

Nashun et al.

Figure S3. Zscan4 is upregulated in the Hira and Cabin1 MII oocytes.

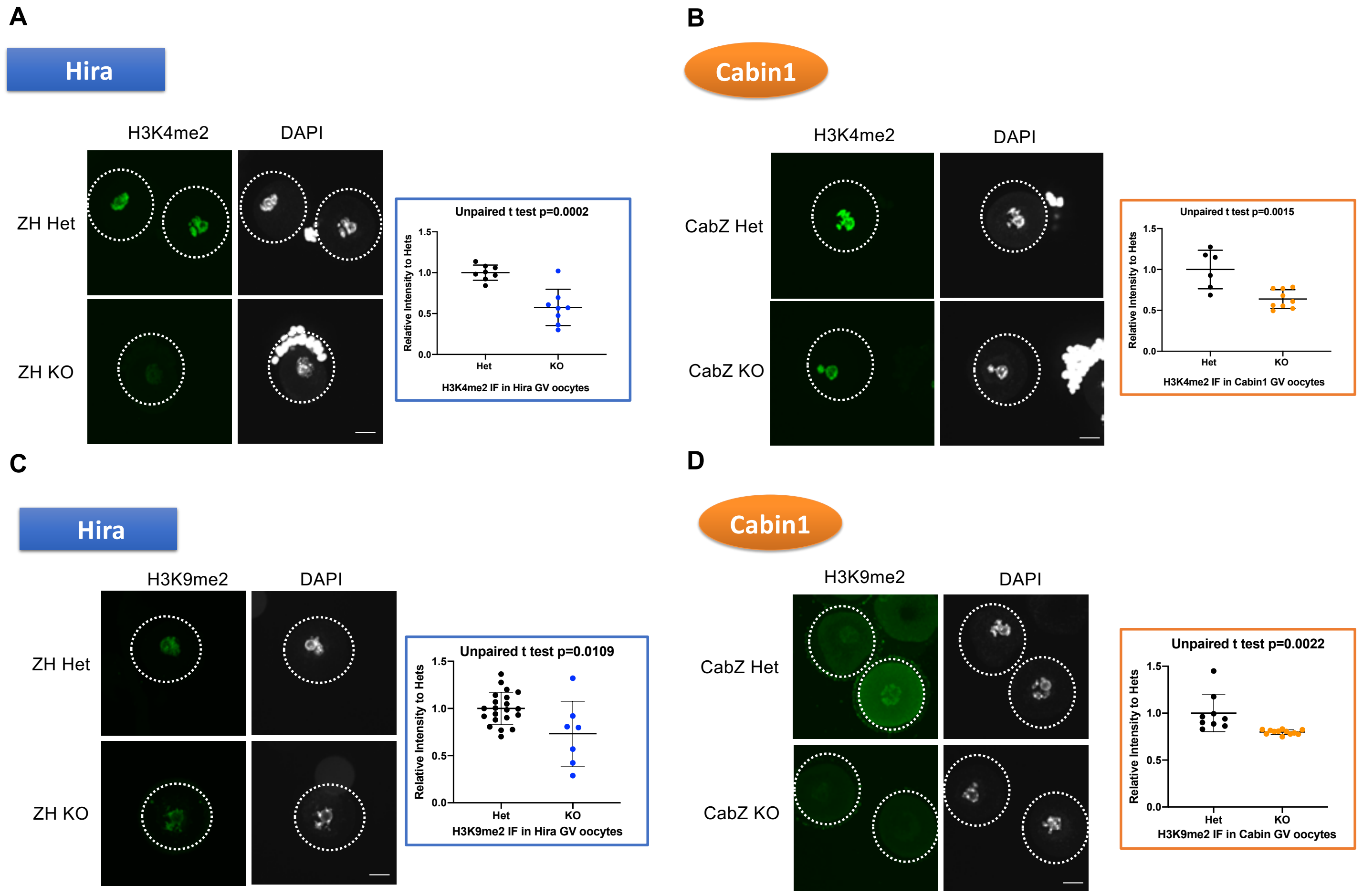


Figure S4-1 Decrease of key histone repressive marks in Hira and Cabin1 mutant oocytes

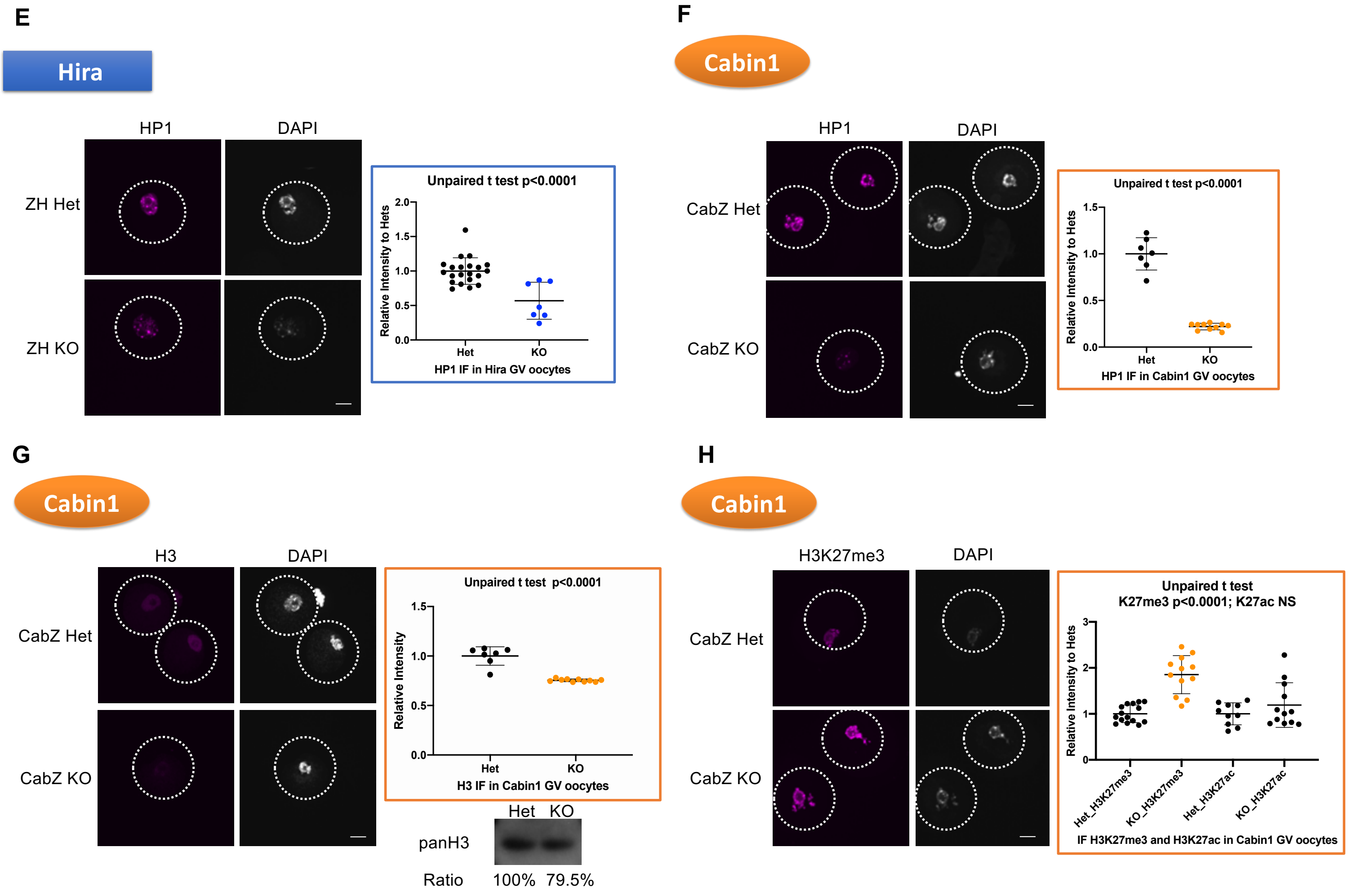


Figure S4-2 Decrease of key histone repressive marks in Hira and Cabin1 mutant oocytes

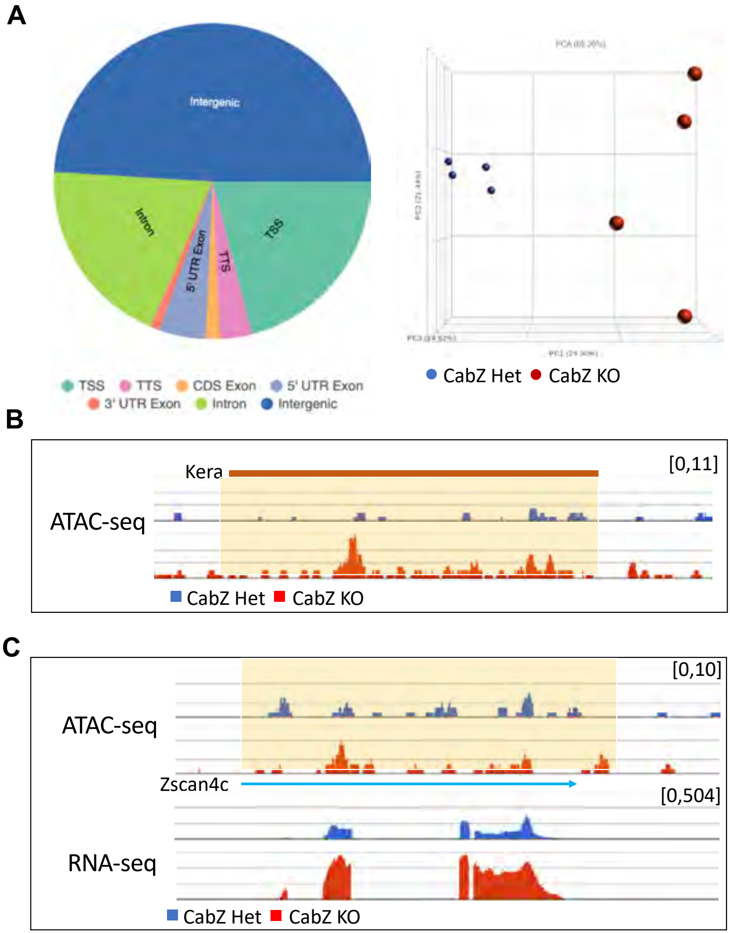


Figure S5. Global chromatin accessibility is increased in Cabin 1mutant GV oocytes

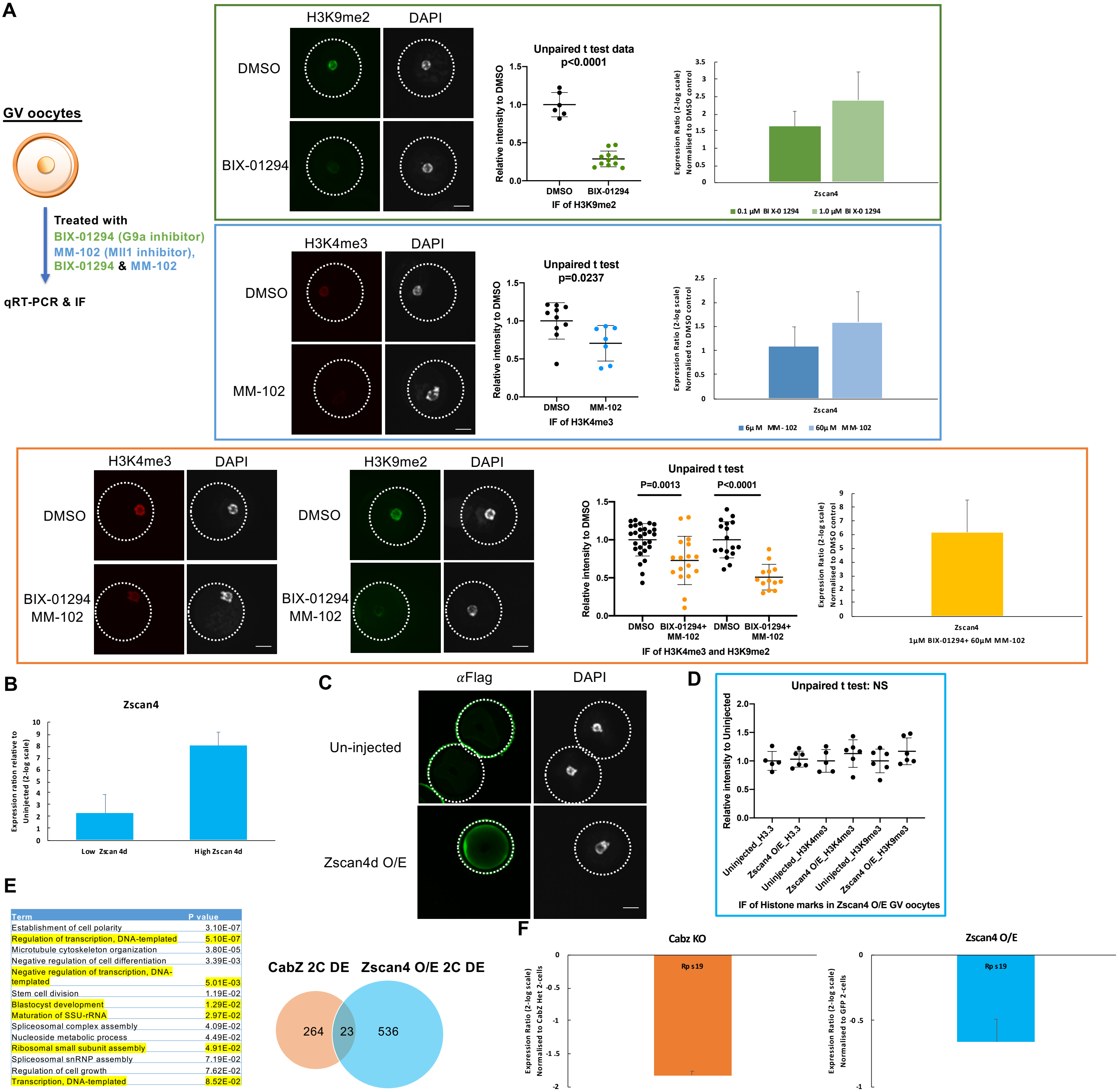


| Term | P value |
|---|---|
| Establishment of cell polarity | 3.10E-07 |
| Regulation of transcription, DNA-templated | 5.10E-07 |
| Microtubule cytoskeleton organization | 3.80E-05 |
| Negative regulation of cell differentiation | 3.39E-03 |
| Negative regulation of transcription, DNA-templated | 5.01E-03 |
| Stem cell division | 1.19E-02 |
| Blastocyst development | 1.29E-02 |
| Maturation of SSU-rRNA | 2.97E-02 |
| Spliceosomal complex assembly | 4.09E-02 |
| Nucleoside metabolic process | 4.49E-02 |
| Ribosomal small subunit assembly | 4.91E-02 |
| Spliceosomal snRNP assembly | 7.19E-02 |
| Regulation of cell growth | 7.62E-02 |
| Transcription, DNA-templated | 8.52E-02 |

Figure S6. Zscan4 expression is regulated by repressive histone marks of H3K4me3 and H3K9me3.

Table S1. List of antibodies used in this study.

| | Manufacturer | Cat. No | Dilution |
|---|---|---|---|
| Flag | Sigma | F3165 | 1:300 |
| Gapdh | Sigma | G9545 | 1:10,000 |
| H3 | Abcam | ab1791 | 1:250 |
| H3K27ac | Abcam | ab177178 | 1:1000 |
| H3K27me3 | Diagenode | C15410195 | 1:200 |
| H3K4me2 | Cosmo Bio | MCA-MABI0003 | 1:1000 |
| H3K4me3 | Millipore | 04-745 | 1:250 |
| H3K9me2 | Abcam | ab1220 | 1:100 |
| H3K9me3 | Abcam | ab8898 | 1:500 |
| HP1 | Cell Signaling | 8676T | 1:250 |
| Laimn A/C | Sigma | SAB 4200236 | 1:500 |
| PolII PS2 | Abcam | 193468 | 1:100 |
| Zscan4 | Gift | Minoru Ko lab | 1:250 |
| Peroxidase Donkey Anti-Rabbit | Jackson Immunoresearch | #711-035-152 | 1:10,000 |
| Alexa Fluor 488 Donkey anti mouse | ThermoFisher | A21202 | 1:500 |
| Alexa Fluor 568 Donkey anti rabbit | ThermoFisher | A10042 | 1:500 |
| Alexa Fluor 647 Donkey anti mouse | Abcam | ab150107 | 1:500 |

Table S2. List of primers used in this study.

| | | |
|---|---|---|
| Zscan4 (d) | Forward | TCCGTAGAGATGCCAAACTATTC |
| | Reverse | GACAGGTGACACAAAGCAATTC |
| Zscan4 (C) | Forward | TTGAAGCCTCCTGTCATGGTCC |
| | Reverse | TCCATTTCATTTCCACTACAGC |
| Ddit3 | Forward | TGTTGAAGATGAGCGGGTG |
| | Reverse | AGGTTCTGCTTTCAGGTGTG |
| Gm5039 | Forward | CCACTGTTAGATACTTCCTGGC |
| | Reverse | AAAAGGTAACCACAGGATCCG |
| Rps19 | Forward | AAGTCCGGGAAGCTGAAAG |
| | Reverse | GAAGCAGCTCGTGTGTAGAA |
